# A single attractor governs the population dynamics of marine fishes worldwide

**DOI:** 10.64898/2026.09.03.749174

**Authors:** Robert M. Hechler, Tianna Peller, Marie-Josée Fortin, Martin Krkosek

## Abstract

Population fluctuations are now empirically recognized to be nonlinear, but the governing attractors are poorly understood. Here we show that population dynamics of 457 marine fish stocks from 121 species worldwide are governed by a single attractor that partitions into three archetypes reflecting top predators, mesopredators and forage fish, with stock-specific ecological and evolutionary variation. Correlation between attractors was high across all stocks (mean 0.53 to 0.63), greater within archetypes (0.57 to 0.77), and increased with geographic, phylogenetic and life history similarity. Leveraging shared attractors increased prediction skill in ∼90% of cases, raising mean recruitment skill from 0.38 to 0.75 and even enabling moderate prediction for otherwise poorly predictable stocks. These results reveal that marine fish population dynamics are not heterogeneous but are realizations of a single attractor structured by trophic level, with immediate predictive benefits for global fisheries.

## Main text

Understanding how population dynamics operate and predicting fluctuations remain fundamental challenges for ecology and management (*1*). Generalizations remain difficult as populations display diverse local adaptations and species differ widely in life histories, environmental conditions and anthropogenic pressures they experience. A dominant paradigm in ecology and evolution assumes that populations fluctuate stochastically around a stable carrying capacity owing to environmental noise (*2–7*). However, a rich body of empirical evidence has accumulated to demonstrate that fluctuations in birds, mammals, insects and marine fishes are instead produced by deterministic nonlinear dynamics, such as oscillations and chaos (*8–16*). Fundamental uncertainties therefore exist whether the rules governing nonlinear dynamics are unique to populations, shared across taxa, or structured by other ecological and evolutionary variables.

The governing rules, or equations, that describe a population’s nonlinear dynamics are encoded in its attractor - a set of points in the phase space that the system converges to, often tracing out shapes such as a fixed point, cycle or more complex geometric patterns (*17–19, 14, 20*). Reconstructing a fully specified attractor is often infeasible because the equations and causal variables are not known a priori. Fortunately, observed population time series are a manifestation of the governing attractor and Takens’ theorem (*21, 22*) enables the use of time lagged copies of these series to reconstruct an empirical attractor that encapsulates the full population dynamics (Figure 1, and see brief introductory video from ref. (*14*) http://tinyurl.com/EDM-intro). Attractor reconstruction methods have been applied broadly across fields such as physics (*21*), climatology (*23*), finance (*24*), epidemiology (*25*), physiology (*26*), genetics (*27*), and in ecology to analyze population dynamics (*8, 9, 11–14*).

**Figure 1.**
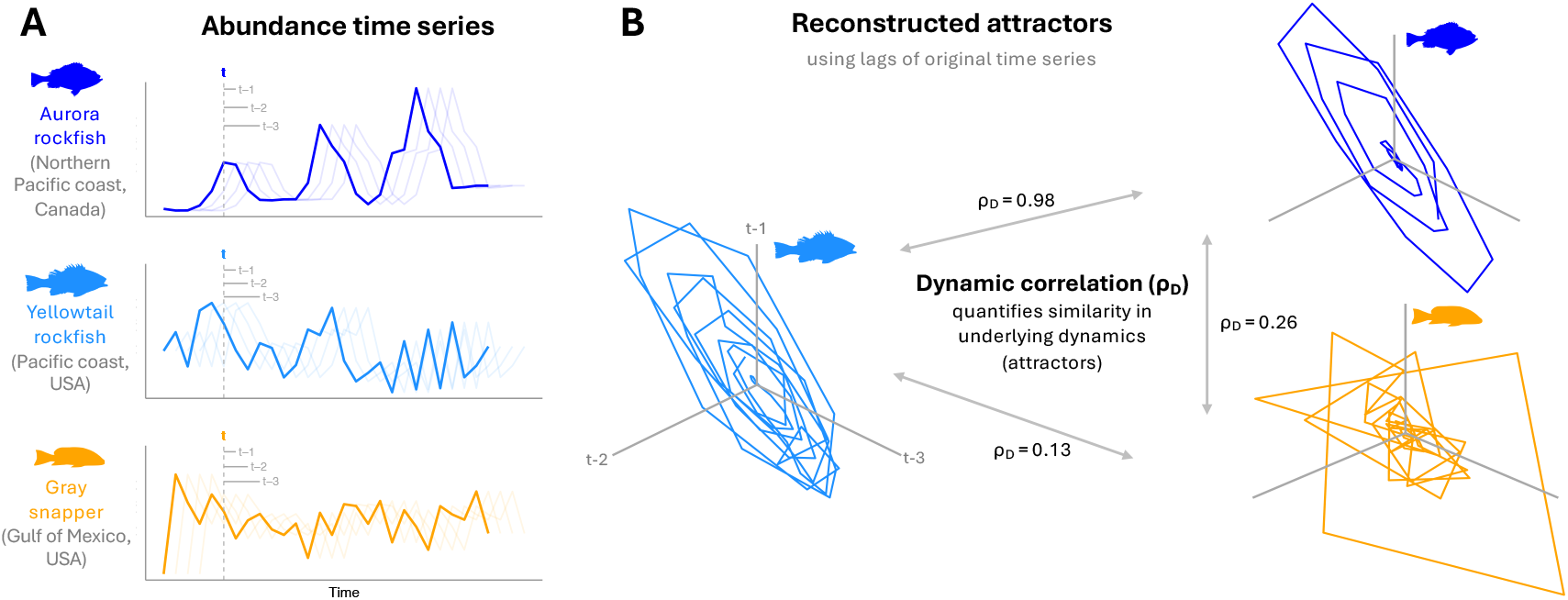
Attractor reconstruction and dynamic correlation. **(A)** Original (dark line) and time-lagged copies (light lines) of recruitment time series for three example species. (**B)** Empirical attractors reconstructed from time lags shown in A. Dynamic correlations (ρ_D_) range between 0 and 1, indicating independent and identical dynamics, respectively. Attractors were reconstructed from smoothed time series for visualization, but all analyses used raw time series.

Several case studies provide evidence that attractors can be similar or shared between populations. Sockeye salmon populations in Canada and Alaska were dynamically correlated despite more than 2000 km of separation, though more weakly than within regions (*28*), indicative of a common species-level attractor that is modified by life-history variation and local conditions. Blue crabs along the US Atlantic coast (*29*) and white shrimp and brown shrimp in the Gulf of Mexico (*30*) similarly had the greatest dynamic correlations among populations in close geographic proximity to one another, providing further support for common species-level attractors shaped by local environmental conditions. Attractors may also extend beyond the species-level, as 23 coastal fish species in California were dynamically equivalent, possibly owing to ecological interactions within the same habitat (*31*). These findings raise the possibility that diverse populations may be organized by common dynamical structures, but the generality, organization and predictive value of such structures remain unknown.

Here, we test the extent to which attractors are shared among marine fish stocks worldwide and whether correlations in the underlying population dynamics are mediated by environmental conditions, life history and phylogeny. We analyzed 218 recruitment and 239 spawning stock biomass time series from the RAM Legacy Stock Assessment Database (*32*) using empirical dynamic modelling (EDM), a non-parametric framework rooted in attractor reconstruction (*33*). Specifically, we used hierarchical EDM to estimate correlation in the underlying population dynamics (dynamic correlation, ρ_D_) (*29, 34*). We then performed a community network analysis using the Louvain algorithm (*35*) to identify archetypes of stocks with similar ρ_D_ values, and multiple regression on distance matrices (*36*) to test whether ρ_D_ dissimilarity was associated with dissimilarity in phylogenetic relatedness, life history traits, geographic distance, and environmental conditions. Lastly, we compared the prediction skill of conventional univariate EDM models that use time lags of the focal stock (Individual model) to multiview EDM models (*37*) that also include information from up to five dynamically correlated stocks, which we refer to as ‘partners’ (Individual + Partners model).

### Shared attractors across species and oceans

The results indicate that population dynamics across species and oceans are governed by a shared attractor structured hierarchically. Network analysis of the top 50% of ρ_D_ identified three groupings in the correlations among recruits and spawners (Fig. 2A,B,D,E). These groupings correspond to archetypes for top predators, mesopredators and forage fish as indicated by differences in PC1 of life history (one-way ANOVA, p = 0.011 for recruits and p < 0.001 for spawners; see Table S1 for pairwise comparisons; see Supplemental Fig. S1 for PCA). Modularity was moderate (0.09 and 0.11; Fig. 3C,D), as expected for a dense network. The groupings are therefore dynamically distinguishable but part of a connected system, indicative of a single community (i.e. group) attractor which all stocks are members of, that partitions into three trophic archetypes. Consistent with this interpretation, ρ_D_ was higher within archetypes than between them, though high levels were shared across all stocks (Fig. 3A,B). Pearson correlations for the fluctuations of stocks within and across archetypes was generally low, despite the high ρ_D_ (Supplemental Fig. S3). The community attractor was reflected qualitatively by consistency in shape and rotation among archetypes (Fig. 3E,F). Within this shared geometry, each archetype has distinct features in size and shape, such as the long barrel for top predators that corresponds to the largest attractor volume (656 for spawners and 444 for recruits; units in standardized state space), followed by mesopredators (382 and 276) and forage fish (245 and 193).

**Figure 2.**
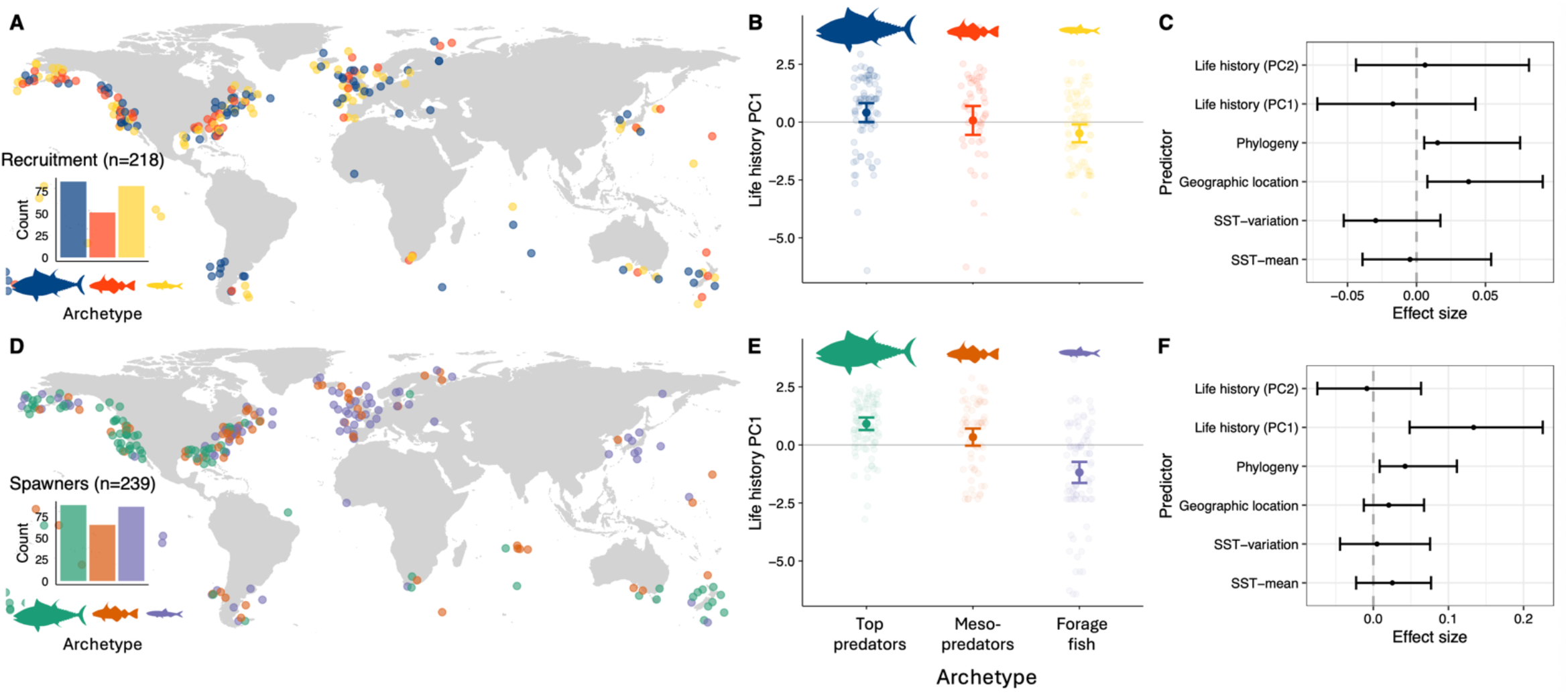
Attractor archetypes and predictors of dynamic correlation. **(A, D)** Spatial distribution of stocks colored by their archetype identified via network analysis (Louvain algorithm) of the top 50% of ρ_D_. See Supplemental Fig. 2 for all ρ_D_. (**B, E)** Archetypes reflect top predator, mesopredator and forage fish groupings as indicated by mean PC1 (error bars are 95% CI; see Supplemental Fig. S1 for PCA). **(C, F)** Predictors of dynamic correlation (dis)similarity. Coefficients were estimated using multiple regression on distance matrices and error bars are bootstrapped 95% confidence intervals (*n* = 1000). SST is sea surface temperature.

**Figure 3.**
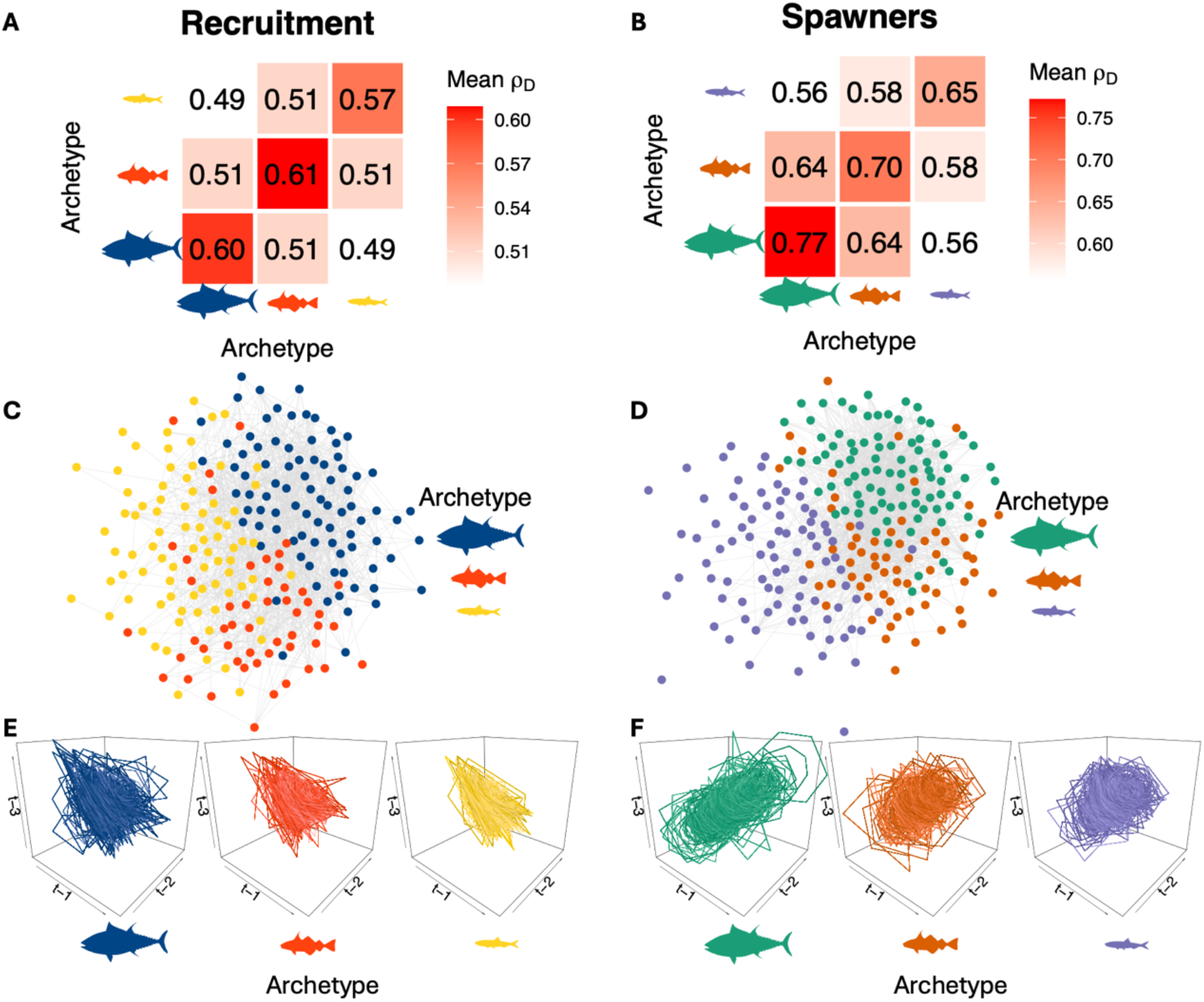
Network structure and attractors of archetypes. **(A, B)** Mean ρ_D_ within and between archetypes. **(C, D)** Network visualized using the Fruchterman-Reingold method. Nodes are stocks colored by archetype. For visual clarity, the strongest 10% of edges (ρ_D_) are shown in grey, but all edges were used to inform the network structure. **(E, F)** Attractors reconstructed using composite time series of all archetype members.

The dynamic correlation across all stocks likely reflects shared foundational processes and membership to a single shared community attractor. Shared dynamics within top predator, mesopredator and forage fish archetypes might reflect processes related to trophic dynamics or their position along the fast-slow life history continuum. These interpretations align with knowledge that marine fish dynamics collapse onto low-dimensional attractors (*11, 12, 38*), with an average embedding dimension (E) of three to four (*12*). E represents the number of variables needed to model the dynamics at a given level of predictability; therefore, despite each population potentially experiencing hundreds of interacting species and abiotic factors, its attractor can be unfolded using only three to four key variables.

### Dynamic correlation increases with geographic, phylogenetic and life history similarity

We used multiple regression on distance matrices (MRM) to test for associations between dissimilarity in ρ_D_ and dissimilarity in life history, phylogeny and environmental factors (Fig. 2C, and 2F). Recruitment ρ_D_ was higher among populations in closer geographic proximity (estimate = 0.038, p = 0.019 and bootstrapped 95% confidence interval (CI): 0.009 to 0.090) and phylogenetic relatedness (estimate = 0.0165 p = 0.256 and 95% CI: 0.005 to 0.072). The other predictors had no clear effects as indicated by 95% CIs that largely overlapped zero. Spawner ρ_D_ was higher among populations with similar PC1 of life history traits (estimate = 0.133, p = 0.0001 and 95% CI: 0.044 to 0.217) and closer phylogenetic relatedness (estimate = 0.042, p = 0.012 and 95% CI: 0.011 to 0.110), while the other predictors had no clear effects. MRM *R*^2^ was 0.0024 for recruitment (F = 9.57, p = 0.13) and 0.013 for spawners (F = 64.11, p = 0.0001), and all p-values were obtained by permutation (9,999 permutations); because MRM operates on pairwise distances rather than raw observations the coefficients and *R*^2^ are systematically lower than conventional regression and should not be interpreted on the same scale (*39–41*).

That ρ_D_ was higher among recruits in close geographic proximity to one another (Fig. 2C) aligns with the emerging view that fluctuations in marine fish stocks result from the amplification of stochastic environmental forcing by nonlinear population dynamics (*10, 12, 13, 42*), the former being more similar among nearby populations. This result generalizes case studies of sockeye salmon (*28*), blue crab (*29*) and brown and white shrimps (*30*) that show greater ρ_D_ among nearby populations. Geographic proximity may be a representation of several abiotic factors that are amplified by the nonlinear skeleton, as similarity in temperature was not associated with ρ_D_ (Fig. 2C,F). This interactive effect of several abiotic variables was shown in Atlantic cod and capelin, where the dynamics were driven by a climate index comprising temperature, sea ice, salinity and seven other abiotic factors (*43*). Similarly, combinations of multiple abiotic variables drove dynamics of sockeye salmon (*12, 44*) and a common reef fish (*13*). The weight of evidence therefore suggests that recruitment dynamics of marine fish arise from the nonlinear amplification of several environmental drivers. However, geographic proximity was not a correlate of spawner ρ_D_, whereas life history (PC1) was strongly correlated (Fig. 2C,F), which accords with known effects of age structure that smooths variation through bet-hedging and portfolio effects (*10, 12, 45, 46*).

The positive associations between ρ_D_ and phylogenetic and life history relatedness (Fig. 2C,F) indicate that the dynamics of functionally similar species are realizations of the same underlying attractor but viewed from different angles. In this framework, the angle at which the attractor is viewed from can be thought of as depending on the realized niche of the population or species (*31*). Thus, just as populations of the same or functionally equivalent species occupy similar niches, they are also likely to display similar dynamics determined by a common attractor, with the viewpoint modified by local environmental conditions. Our results and interpretation are supported by an analysis of the Georges Bank fish community in northeastern USA, where 26 species clustered into four groups of dynamically equivalent species reflecting similar feeding patterns and habitat use (*47*). The same attractor could in principle be recovered from different rules, for example if it was a simple point attractor representing a stable equilibrium. But marine fishes are notorious for their nonlinear fluctuations (*10–13, 15, 16*), and the consistent shape and ordering of attractor size across trophic levels would be unlikely produced by unrelated dynamics.

### Shared attractors enable improved prediction

If time series of different stocks are indeed produced by the same underlying attractor as indicated by Figs. 2 and 3, including information from dynamically correlated stocks (which we refer to as ‘partners’) should provide a more complete view of the shared attractor and therefore increase prediction skill. As expected, prediction skill improved substantially when using dynamically correlated partners to reconstruct different views of the shared attractor, where *k* is the number of reconstructed views (Fig. 4 and Supplemental Fig. S4). Mean recruitment prediction skill of the Individual model was 0.38 (standard deviation, s.d. = 0.27) and increased when ensembling all *k* views to 0.65 (s.d. = 0.17) and to 0.75 (s.d. = 0.12) when ensembling the top-ranked 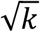 views (Fig. 4A,B). Mean spawner prediction skill using the Individual model was 0.77 (s.d. = 0.22) and increased to 0.91 (s.d. = 0.09) when ensembling all *k* views and to 0.95 (s.d. = 0.05) when ensembling the top-ranked 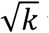 views (Fig. 4D,E). Leveraging information from partners was most beneficial for stocks that were poorly predictable using their own time lags. For recruitment stocks in the lower 50th percentile, mean Individual prediction skill was 0.16 (s.d. = 0.17) but increased to 0.59 (s.d. = 0.15) and 0.71 (s.d. = 0.11) when ensembling all *k* and the top 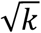 views, respectively (Fig. 4C). For spawners, mean Individual prediction skill was 0.62 (s.d. = 0.22) and increased to 0.87 (s.d. = 0.09) and 0.92 (s.d. = 0.06), respectively (Fig. 4F).

**Figure 4.**
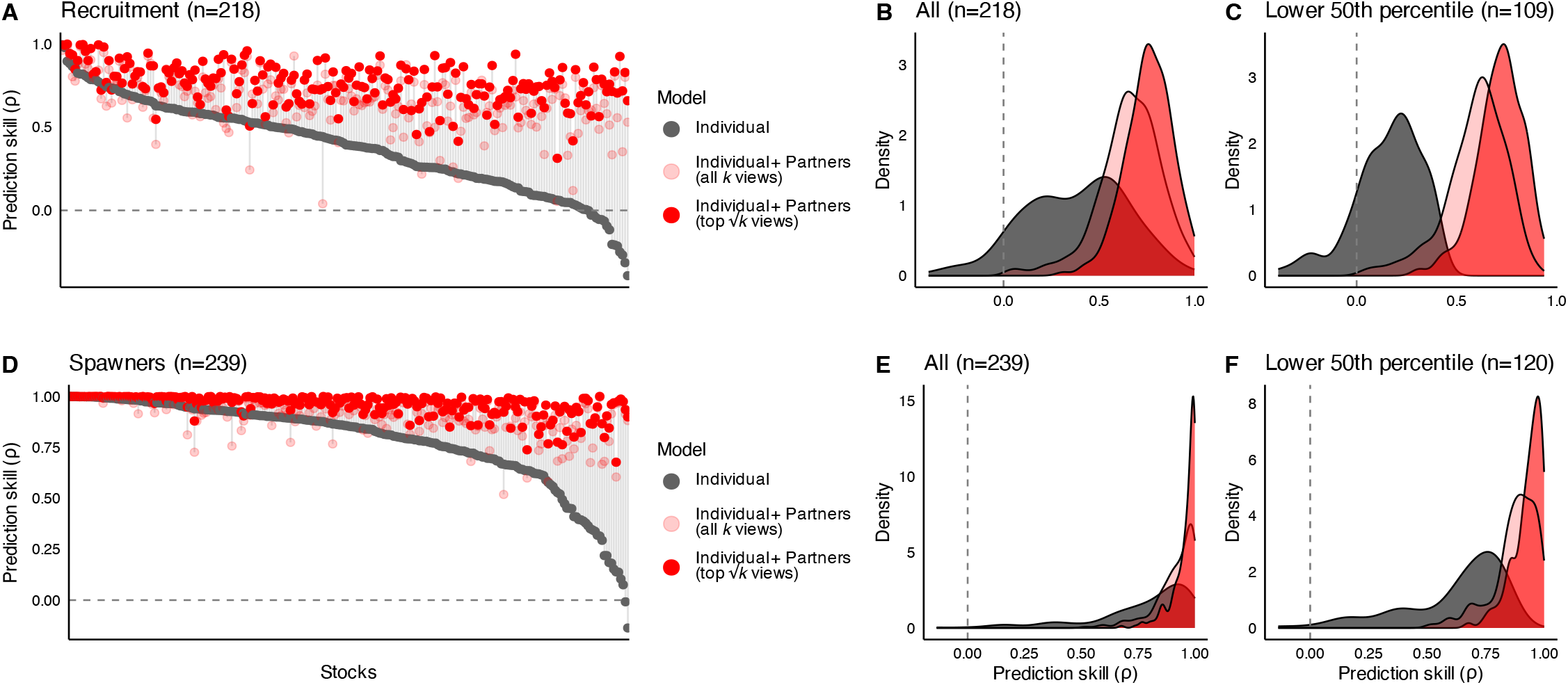
Leveraging shared attractors improves prediction skill. **(A, D)** Prediction skill (Pearson correlation between observed and predicted values) for two models: *Individual*, univariate EDM using only the focal stock’s own lags (grey) and *Individual + Partners*, multiview EDM including also information from up to five dynamically correlated partner stocks (red). Each three-dimensional combination of focal and partner lags is a view of the attractor. **(B, E)** Density plot of prediction skill across all stocks. **(C, F)** Density plot of prediction skill for stocks in the lower 50th percentile of the Individual model.

The substantial and systematic gain in prediction skill by leveraging dynamically correlated partner stocks provides further evidence for shared attractors across fish stocks. Individual + Partners models outperformed the Individual model for 97% (top 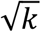 views) and 89% (all *k* views) of recruitment stocks and for spawners in 98% and 85% of stocks. This result is especially promising given that accurate prediction is essential for management but remains difficult in practice, with two meta-analyses of 185 and 211 fish stocks showing standard parametric fisheries models (Ricker and Beverton-Holt) to explain only 4 to 15% of variance in recruitment (*16, 48*). Although non-parametric EDM methods are already known to outperform forecasts from parametric fisheries models (*16, 44*), many stocks remained poorly predictable using individual EDM in a meta-analysis (*16*) and here (Fig. 4A). Leveraging shared attractors from stocks that are phylogenetically similar and in close geographic proximity improves recruitment forecasts and could therefore aid management.

Overall, our study provides evidence that the population dynamics of the world’s marine fish stocks are governed by a single community attractor that partitions into three archetypes of top predators, mesopredators and forage fish, within which stocks vary according to ecological and evolutionary factors. Correlation in the underlying dynamics increased for recruits with geographic proximity and phylogenetic similarity, while for spawners it increased with phylogenetic and life history similarity. Leveraging information from dynamically correlated stocks improved prediction skill systematically, even enabling moderate prediction of stocks that were otherwise poorly predictable. These results suggest that shared attractors are a general feature of marine fish stocks across the world, with immediate predictive benefits for fisheries, conservation and management.

## Materials and Methods

### Data

The original fish abundance and biomass time series data are from the RAM Legacy Stock Assessment Database (RAM LSAD) (*32*), stock boundaries from ref. (*49*), sea surface temperature from the National Oceanic and Atmospheric Administration Optimum Interpolation SST V2 High Resolution database (*50*), life history traits from fishLife (*51*) and a time-calibrated phylogeny (*52*). We previously compiled and cleaned these data in ref. (*12*). Spawner data represent estimates of adult biomass, while recruitment represents estimates of the number of young individuals that survive early periods of density-dependent mortality and subsequently ‘recruit’ into the population.

### Network analysis and multiple regression on distance matrices

Dynamic correlation (ρ_D_) between each pair of stocks was estimated by fitting hierarchical Gaussian process empirical dynamic models (GP-EDM) models (*29, 34*) with a maximum embedding dimension (E) of 10. We then performed network analyses of ρ_D_ via the Louvain algorithm to identify attractor archetypes. As the Louvain algorithm implements stochastic node ordering, we performed the analysis 100 times and retained the modal archetype count and median modularity score. We performed this network analysis on the top 50% of ρ_D_ (Fig. 2), as well as on all ρ_D_ (Supplemental Fig. S2).

We performed multiple regression on distance matrices (9,999 permutations) to test whether dissimilarity in ecological and evolutionary factors is associated with dissimilarity in ρ_D_. The response was ρ_D_ dissimilarity (1-ρ_D_) and predictors were pairwise distances for geographic location (centroid of stock boundary), differences in mean sea surface temperature (SST), intra-annual SST variation, phylogenetic distance and two principal components of life history traits (Supplemental Fig. S1). Bootstrapping stocks with replacement was used to calculate 95% confidence intervals (*n* = 1000). Predictor matrices were normalized between 0 and 1 for comparability.

### Predicting recruit and spawner stocks

As a baseline prediction skill to compare against, we fit an “Individual” GP-EDM model in the conventional univariate setting using only lags of the target stock as predictors. The embedding dimension (E) was set to the square root of the time series length (with a maximum E of 10), the time lag was one, and prediction skill was measured as the Pearson correlation between observed and predicted values. All models were fit in R version 4.5.1 (*53*) using the GP-EDM R package (*54*) and predictions were always made out of sample via leave-one-out-cross validation with an exclusion radius equal to E.

We used multiview embedding with up to five dynamically correlated stocks (which we refer to as partners) to test whether leveraging information from shared attractors improves prediction skill. For each focal stock, we identified partners by testing how well the focal stock was predicted by each of its 50 most dynamically correlated stocks. A separate model was fit per candidate using lags 0 to 4 of that candidate as predictors and no lags of the focal stock. The five stocks with the highest prediction skill were selected as partners and for each, we retained the lag with the greatest inverse length scale for use in multiview embedding. Multiview embedding reconstructs a system’s attractor using several different low-dimensional combinations of variables (*37*). The idea is that a common attractor is shared between the focal and partner stocks, and each combination of their time lags represents an equally valid reconstruction of that shared attractor but viewed from different angles (*37*). Thus, averaging predictions made across each unique view (*k*) should reduce noise and improve prediction skill. We reconstructed three-dimensional attractors from different combinations of the focal’s four lags and one lag per partner, and report prediction skill for ensembles of all *k* views as well as ensembles of the top 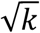 views. We also performed a sensitivity analysis restricting the partner search only over the 10 most dynamically correlated stocks (rather than 50) and the results were qualitatively similar (Supplemental Fig. 4).

## Author contributions

R.M.H. and M.K. designed research; R.M.H. led the analysis with input from M.K., M-J.F. and T.P.; R.M.H. wrote the first draft of the manuscript and M.K., M-J.F. and T.P. contributed substantially to revisions.

## Code availability

All code and analyzed data will be made publicly available upon manuscript acceptance.

