## Supplementary information for "A single attractor governs the population dynamics of marine fishes worldwide"

**Table S1.** Pairwise comparisons of mean PC1 values among archetypes from Tukey's HSD (Honest Significant Differences) tests. 1, 2, and 3 represent top predator, mesopredator and forage fish archetypes, respectively.

| **Data type** | **Comparison of archetypes** | **Difference** | **Lower 95 CI** | **Upper 95 CI** | **Adjusted p-value** |
|---|---|---|---|---|---|
| Spawner biomass | 2-1 | -0.574 | -1.222 | 0.074 | 0.094 |
| Spawner biomass | 3-1 | -2.095 | -2.696 | -1.494 | <0.001 |
| Spawner biomass | 3-2 | -1.521 | -2.173 | -0.870 | <0.001 |
| Recruitment | 2-1 | -0.339 | -1.143 | 0.465 | 0.580 |
| Recruitment | 3-1 | -0.895 | -1.60 | -0.191 | 0.008 |
| Recruitment | 3-2 | -0.556 | -1.369 | 0.257 | 0.242 |

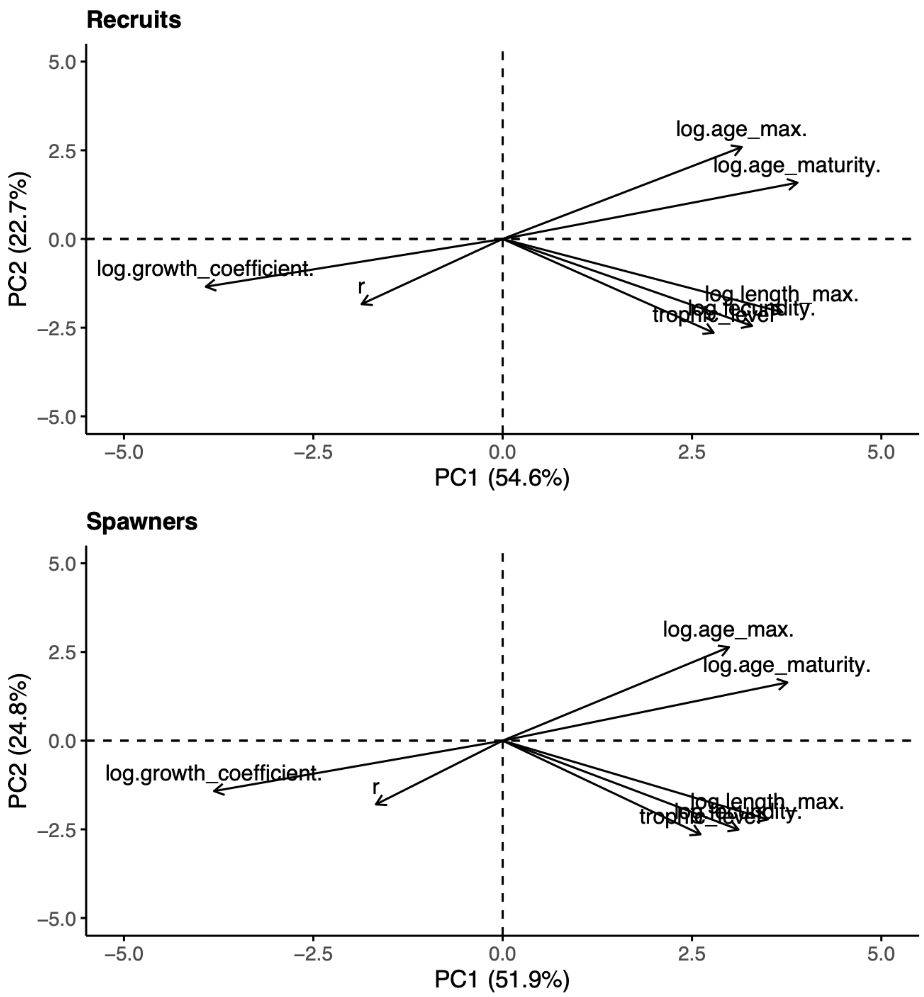


Supplemental Figure S1. Principal component analysis (n = 218 recruitment and 238 spawner stocks) of the following species-level life history traits: trophic level, intrinsic growth rate (r), log-growth coefficient, log-max age, log-max length, log-age at maturity and log-fecundity.

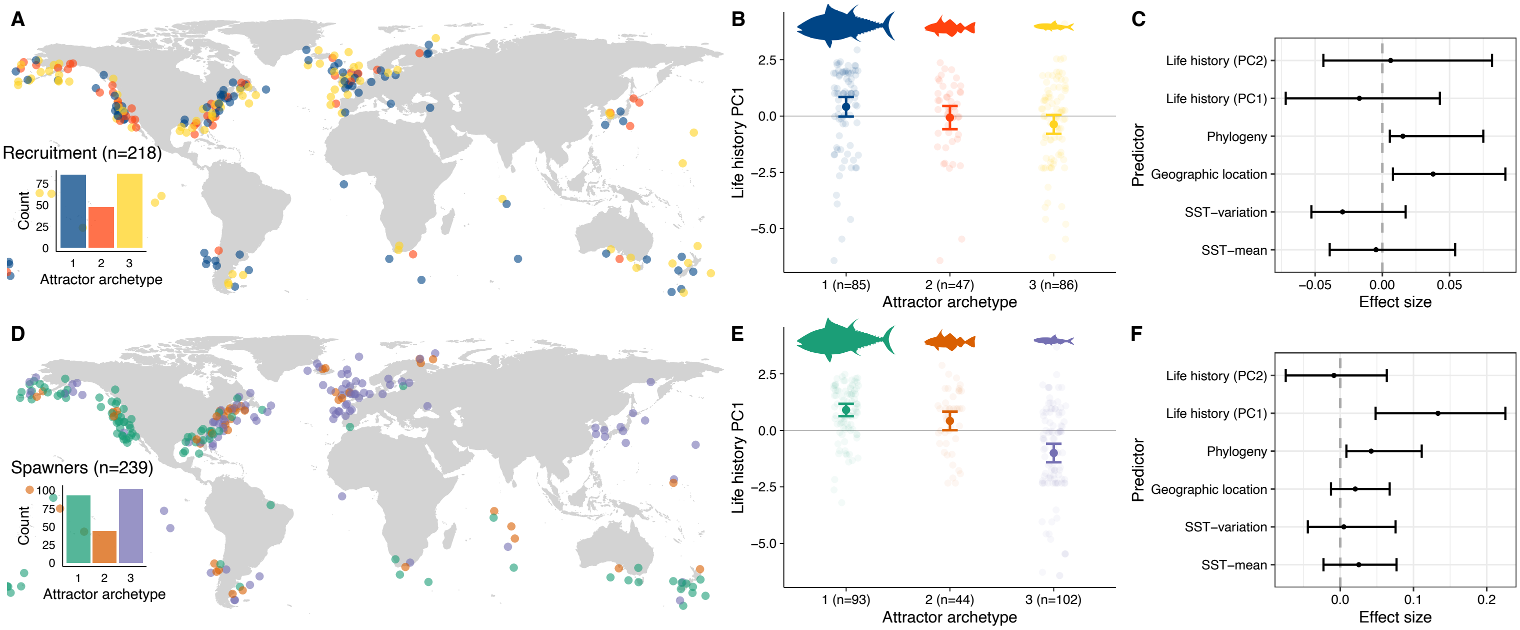


Supplemental Figure S2. Same as Figure 2 but using all dynamic correlations in the network analysis instead of the top 50%. 1, 2, and 3 represent top predator, mesopredator and forage fish archetypes, respectively. Panels C and F are identical to Figure 2 C and F.

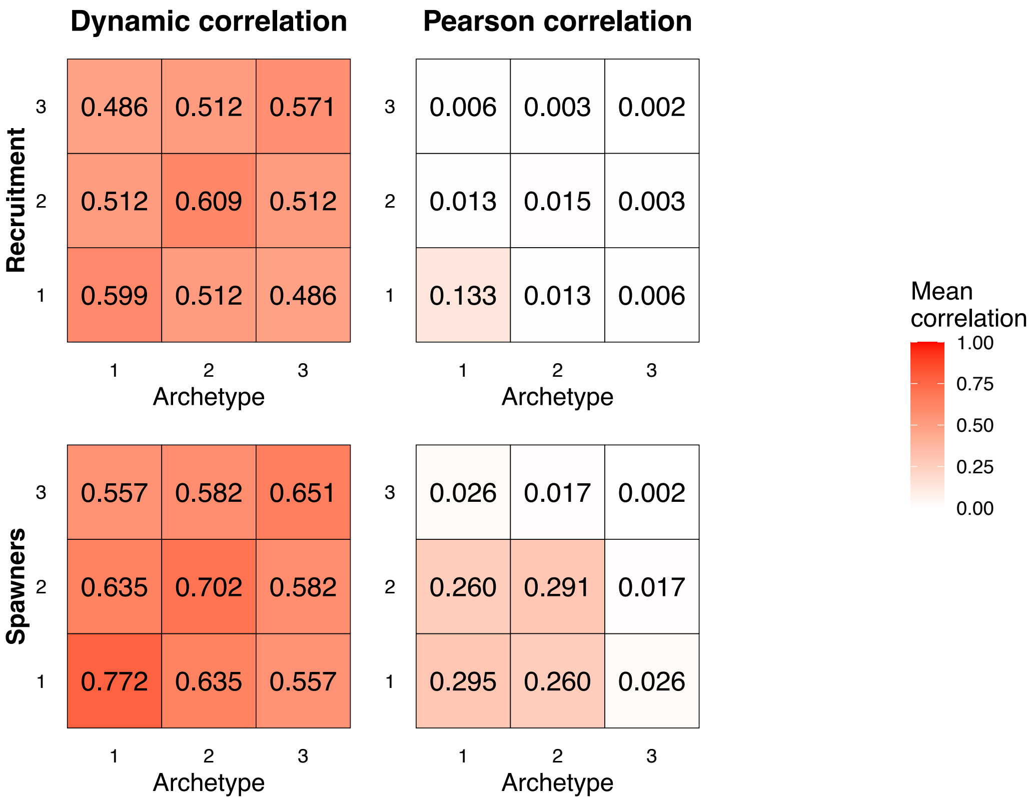


Supplemental Figure S3. Fluctuations are uncorrelated or weakly correlated despite strong correlation in the underlying population dynamics as indicated by mean dynamic and Pearson correlations of stocks within and across archetypes. Top row is recruitment and bottom is spawners. 1, 2, and 3 represent top predator, mesopredator and forage fish archetypes, respectively.

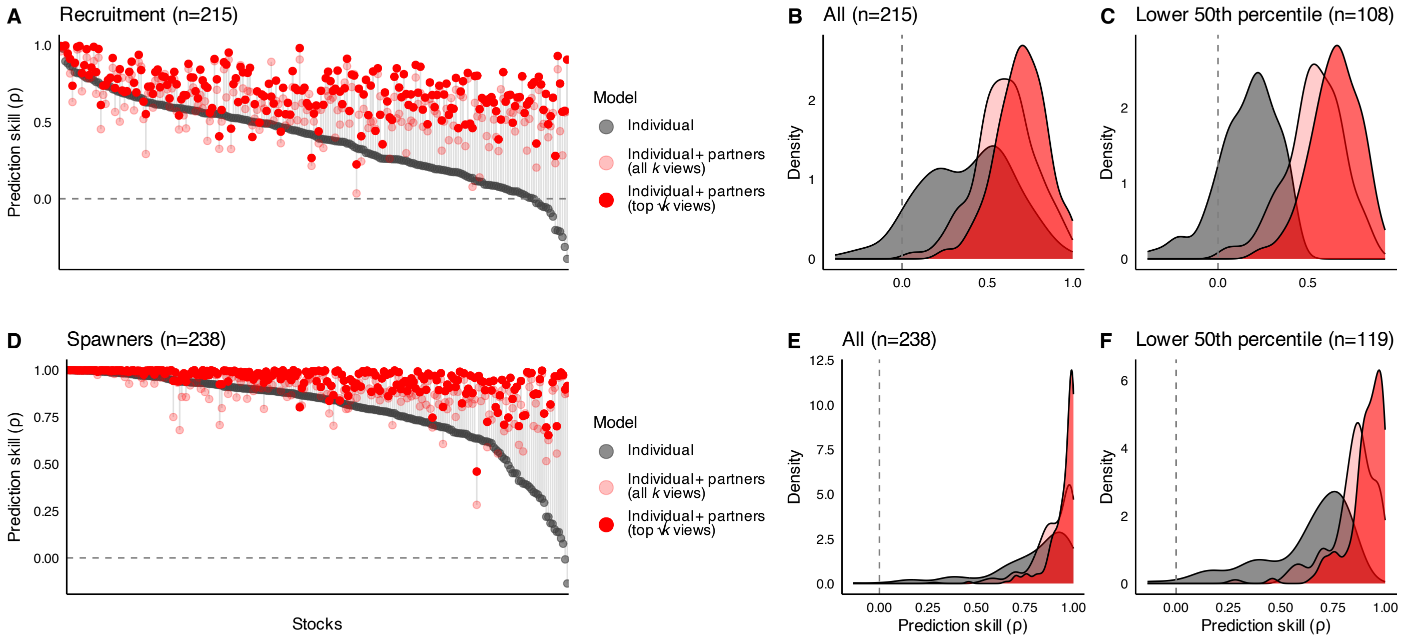


Supplemental Figure S4. Same as Figure 4 but testing only the 10 most dynamically correlated stocks as potential ‘partners’ (rather than 50). Three recruitment and one spawner stock were removed as all ten potential partner stocks had negative predictive skill.
